# Diversity decoupled from sulfur isotope fractionation in a sulfate reducing microbial community

**DOI:** 10.1101/518837

**Authors:** Jesse Colangelo, Claus Pelikan, Craig W. Herbold, Ianina Altshuler, Alexander Loy, Lyle G. Whyte, Boswell A. Wing

## Abstract

The extent of fractionation of sulfur isotopes by sulfate reducing microbes is dictated by genomic and environmental factors. A greater understanding of species-specific fractionations may better inform interpretation of sulfur isotopes preserved in the rock record. To examine whether gene diversity influences net isotopic fractionation *in situ*, we assessed environmental chemistry, sulfate reduction rates, diversity of putative sulfur metabolizing organisms by 16S *rRNA* and dissimilatory sulfite reductase (*dsrB*) gene amplicon sequencing, and net fractionation of sulfur isotopes along a sediment transect of a hypersaline Arctic spring. *In situ* sulfate reduction rates yielded minimum cell-specific sulfate reduction rates <0.3 x 10^−15^ moles cell^−1^ day^−1^. Neither 16S *rRNA* nor *dsrB* diversity indices correlated with relatively constant (38 to 45‰) net isotope fractionation (ε^34^S_sulfide−sulfate_). Measured ε^34^S values could be reproduced in a mechanistic fractionation model if 1-2% of the microbial community (10-60% of Deltaproteobacteria) were engaged in sulfate respiration, indicating heterogeneous respiratory activity within sulfate-metabolizing populations. This model indicated enzymatic kinetic diversity of Apr was more likely to correlate with sulfur fractionation than DsrB. We propose that, above a threshold alpha diversity value, the influence of the specific composition of the microbial community responsible for generating an isotope signal is overprinted by the control exerted by environmental variables on microbial physiology.

**Subject categories:**

i. Integrated genomics and post-genomics approaches in microbial ecology
ii. Microbial ecology and functional diversity of natural habitats

## 1. Introduction

During microbial sulfate reduction (MSR), lighter isotopologues of sulfur are reduced at a greater rate than heavier ones, resulting in sulfide isotopically enriched in light isotopes of sulfur relative to the source sulfate (1). Variations in biologically mediated fractionation result from environmental differences including sulfate concentration, electron donor type and concentration, and temperature (2–9). These variables influence cell-specific sulfate reduction rate (csSRR), and a frequently observed relationship between csSRR and sulfur isotope fractionation (Fig. 1A) has been used to interpret the environmental conditions associated with the time and place of sulfate reduction (10). Tracking biological fractionation of sulfur isotopes through interpretation of mineralized sulfur compounds provides a metric of biogeochemical activity and the role of microbial metabolisms in the cycling of sulfur, both in modern and ancient environments (11). Strikingly, measured differences in environmental sulfur isotope fractionation are only ever interpreted in terms of environmental controls, despite strong evidence for species-specific isotope enrichment effects (Fig. 1B)(12–14). There is no known relationship between microbial diversity and isotopic fractionation. This potentially confounds environmental interpretations of the sulfur isotope rock record, given the multitude of contemporary microbes capable of MSR and the lack of techniques to establish which, if any, of these genotypes were present and active at the time an isotope fractionation signal was generated. In the last decade, research has shifted towards exploring species-specific physiological contributions to the extent of isotope fractionation (15–17), but studies approaching the problem from the opposite direction, interrogating entire communities, are lacking.

**Fig. 1.**
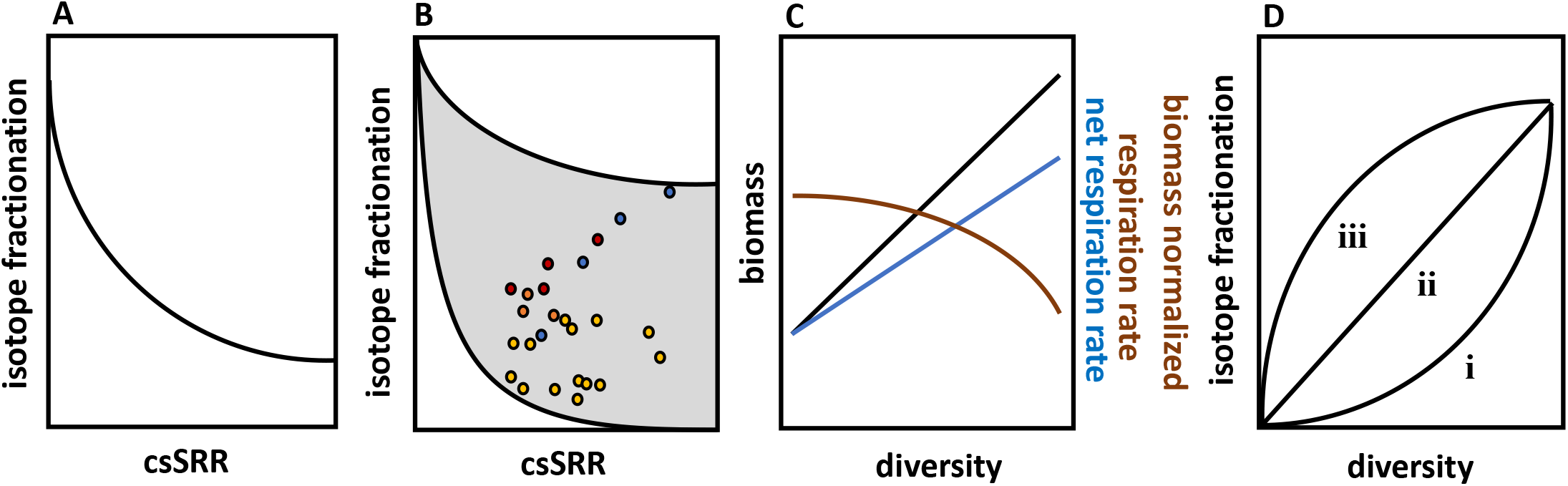
Experimental rationale. A) This schematic representation of data presented in Harrison and Thode 1958, Kaplan and Rittenberg 1964 and Chambers *et al*. 1975 illustrates the relationship between cell-specific sulfate reduction rate (csSRR) and the extent of isotope fractionation that has been used to interpret environmental conditions throughout Earth’s history. B) Grown under the same conditions, different species of sulfate reducing bacteria fractionate isotopes of sulfur to widely varying extents, demonstrating the relationship in A) is species-specific. Each data point represents a different sulfate reducing species and each color represents a distinct set of growth conditions (modified from Detmers et al. 2001). This data set argues for a wide range of fractionation-csSRR trajectories (shaded), bound to an upper maximum by equilibrium fraction. C) Ecological theory predicts environments with greater diversity will maintain a greater biomass and a greater net rate of respiration (e.g. Cardinale *et al*., 2006), though biomass-normalized respiration rate may decrease if increasing species are functionally redundant (e.g. Power and Cardinale, 2009). D) Given a negative relationship between fractionation and csSRR, and a negative relationship between biomass-normalized respiration (comparable to csSRR) and diversity, we expect a positive relationship between isotope fractionation and diversity. The trajectory of this relationship is unconstrained by this hypothesis. Trajectory i represents a lag in metabolic redundancy with increasing diversity and progressively more redundancy with more diversity, trajectory ii represents a direct relationship between metabolic redundancy and diversity, trajectory iii represents a rapid increase in redundancy with increasing diversity and diminishing redundancy after an inflection point with diversity.

Experimental work has confirmed the prediction from ecological theory that environments with greater species richness will maintain a greater biomass and exhibit a greater net rate of respiration (i.e. carbon respired per time (Fig. 1C)(18,19). However, while net respiration increases with species richness, respiration per biomass in more species-rich environments can decrease if species exhibit functional redundancy (Fig. 1C)(20,21). This is relevant to our system because of the inverse relationship between csSRR and isotope fractionation (Fig. 1A)(1). More species-rich environments typically bear a greater complexity of organic matter, and increasing complexity of organic matter utilized in sulfate respiration leads to lower csSRR and higher fractionation (22). Given the greater extent of fractionation exhibited by SRM reducing sulfate at low rates, and the greater influence on net fractionation by these organisms (Fig. 1A), we predicted communities with a greater diversity to be positively correlated with net fractionation of sulfur isotopes, irrespective of csSRR within that environment, though the shape of that trajectory is expected vary based on the nature of metabolic redundancy (Fig. 1D).

Our prediction rested on two premises: First, increasing diversity of an environment does not favor organisms with greater csSRR within that environment (which would drive net fractionation to lower values) and second, when the diversity of an environment increases, SRM with lesser csSRR reduce a greater number of sulfate molecules than SRM with a greater csSRR compared to the previous population. Both of these premises are illustrated by the equation below where [SRM] is the abundance of any new SRM contributing to an increase in community species richness, ε is the species-specific fractionation of sulfur isotopes and csSRR is the cell-specific rate of sulfate reduction.

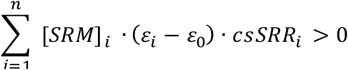

These premises are untested and results that do not support our prediction might be attributed to one or both of them.

The goal of this study was to investigate the role of a microbial diversity, in a natural setting, on the net fractionation of sulfur isotopes. A sediment transect from a hypersaline Arctic spring with perennial and consistent geochemistry (23) was chosen to explore these questions. This spring harbors a known low diversity of SRM where individual contributions to net fractionation by a given gene type are expected to be more pronounced (24). We collected geochemical data along a transect extending from the spring’s outlet to 20 m downstream. From the same sediments, we measured *in situ* SRRs, multiple isotopes of sulfur, and SRM diversity. We sequenced the canonical microbial diversity 16S rRNA gene, as well as the gene *dsrB* that encodes the beta-subunit of the SRM key enzyme dissimilatory sulfite reductase (25), which catalyzes initial steps in the reduction of sulfite to sulfide (16,26). The DsrAB enzyme cleaves three of the four S-O bonds of the respired sulfate molecule and has been proposed to act as an isotopic bottleneck in sulfate reduction, suggesting changes in the diversity of the gene *dsrB* might disproportionately influence fractionation (1,27).

## 2. Methods

### Site description

Samples were collected from the largest of the Gypsum Hill springs (spring GH4) on Axel Heiberg Island, Nunavut (Fig. 2). The geochemistry of the spring over more than a decade has been relatively constant (23). The spring’s outlet water is perennially cold (4.2-6.9 °C), circumneutral pH (7.4), hypersaline (75-82 ‰), near anoxic (<6 μM dissolved oxygen), and reducing (−320 to −285 mV). Microbial culturing and molecular characterization of the spring outlet sediments have identified bacterial and archaeal taxa and genes associated with aerobic and anaerobic heterotrophic and autotrophic metabolisms, including sulfur and sulfate reducing bacteria, methanogens and methanotrophs (24,28,29). Samples and measurements for this study were taken from the spring outlet (Outlet, 0 m) and from channel stations at 1, 4, 8, 12, and 18 m downstream (Channel 1-5).

**Fig. 2.**
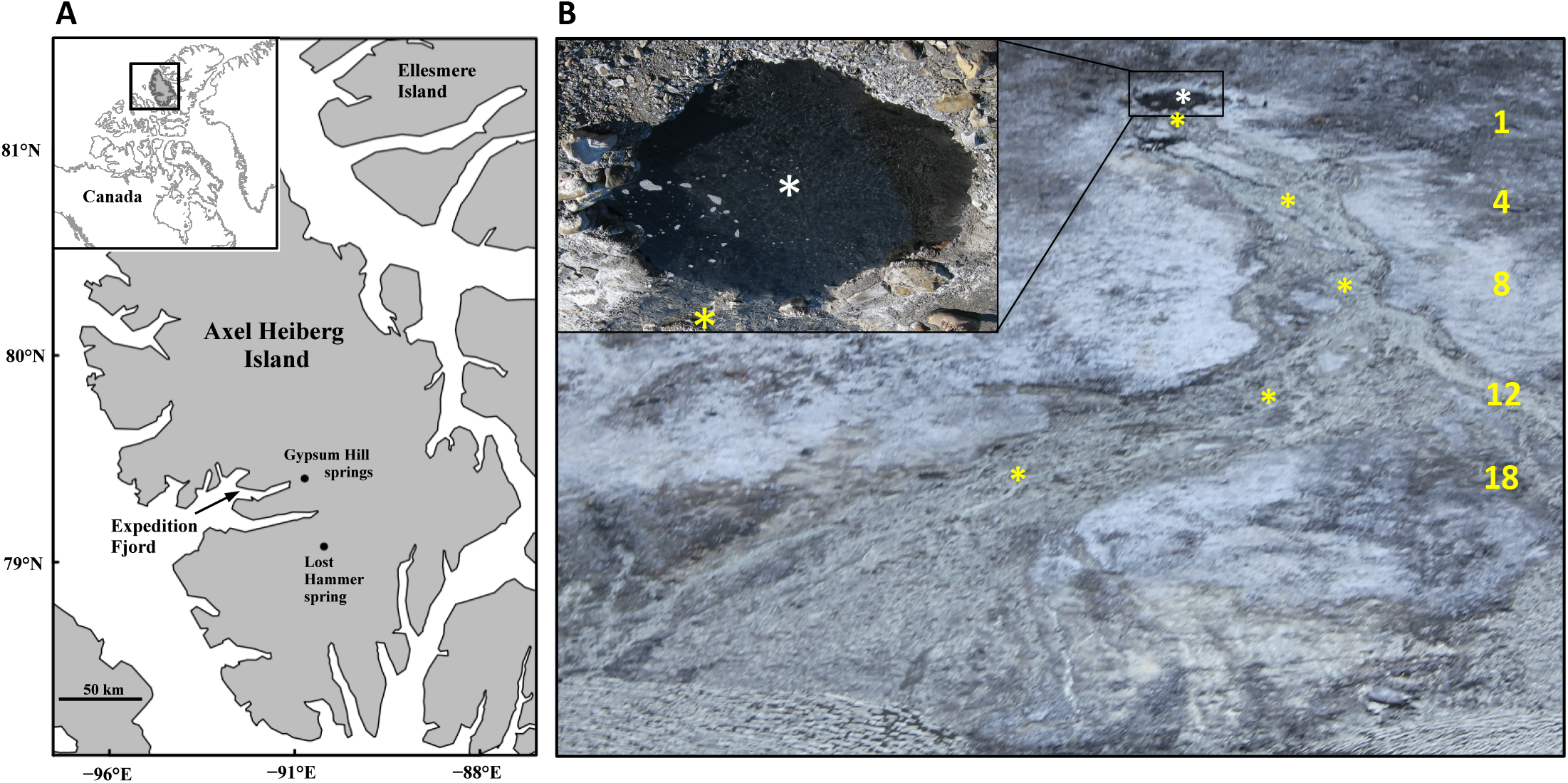
Gypsum Hill Spring: location and sampling stations. A) Gypsum Hill Spring is located on Axel Heiberg Island in the Canadian High Arctic. B) Sediment collection stations are indicated by asterisks (white: Outlet; yellow: Channel). Sampling station distances from Outlet, along flow path, are noted. Outlet pool (inset) is ~2 m in diameter. Distance (m) from outlet pool to spring terminus (Expedition River, bottom left) is ~ 30 m.

### Sediment and pore water measurements

Pore water measurements were taken at 3 cm sediment depth from each sampling station and included temperature, pH, and concentrations of dissolved oxygen, sulfide, and organic carbon. Oxidation-reduction potential, concentrations of pore water nitrate, nitrite, ammonia, thiosulfate, ferric iron, phosphate, and sediment total carbon and nitrogen were assessed from the outlet and a single channel station, 8 m downstream of the outlet (Channel-3). For chemical analyses, sediment pore water was collected into syringes through microfiltration (0.12 μm) membrane tubing inserted into the sediment and assessed for dissolved ions immediately upon recovery. Pore water samples for dissolved organic carbon measurement were filtered through GF/F filters, acidified and transported in precombusted glass vials. Sediments for solid phase carbon and nitrogen were collected with a sterile spatula into polypropylene tubes. Following acid conversion of inorganic carbonates to carbon dioxide, total organic carbon and nitrogen were measured at the GEOTOP Stable Isotope Laboratory at UQAM (Montreal, QC). Dissolved organic carbon was measured at the GEOTOP Environmental Organic Geochemistry Laboratory at Concordia (Montreal, QC). Sediment porosity was calculated from sediment density and mass reduction following drying at 100°C for 24 h. Specific instruments and kits are listed in Supplemental Materials Table S1.

### Sediment collection for sulfur isotopes and DNA

Sediments for isotope and DNA analyses were collected as described above. Spring water and sediment samples for analysis of sulfur isotopes were treated with 20 % wt/vol acidified Zinc acetate (ZnAce), and stored and transported frozen. Additional spring water samples were treated with BaCl2 to precipitate aqueous sulfate. Sediments for DNA extraction were stabilized with RNALifeguard and stored and transported at −5°C.

### Sulfur isotope composition of spring water and sediments

Mineralized forms of various oxidation states of sulfur were serially extracted from 5 g wet sediment for isotopic analysis. Water-soluble sulfate was precipitated from extracted sediments with BaCl_2_ and reduced to sulfide by reacting 10 mg of BaSO_4_ with 15 mL of Thode solution (32:15: HI:H_3_PO_2_:HCl) at 100°C under a stream of N_2_ for 90 min (30). N_2_ carried generated H_2_S to a ZnAce trap to quantitatively precipitate H_2_S as ZnS. Excess AgNO_3_ (0.1 N) was added to the ZnAce traps to convert ZnS to Ag_2_S. Ag_2_S was separated from ZnAce by filtration on 0.2 μm nitrocellulose filters, rinsed with ammonium hydroxide and then MilliQ water, scraped from the filters, and dried overnight at 50°C. The same distillation apparatus and conversion to Ag_2_S was employed in each serial extraction. Sediments were then extracted with MeOH to solubilize elemental sulfur (S^0^). MeOH was decanted and evaporated. Residual S^0^ was converted to H_2_S by hot HCl distillation in the presence of 1M Cr^2+^. MeOH-extracted sediments were washed twice in MilliQ water. Acid volatile sulfides (AVS) were extracted with hot HCl and precipitated as above. Following collection of trapped ZnS from the acid distillation, and without removal from the distillation apparatus, sediments were extracted for chromium reducible sulfides (CRS) as above. Both decanted supernatant from this acidic chromium extraction, as well as residual sediments were separately extracted with Thode solution in the same manner as the BaSO_4_ precipitated above. Ag_2_S from all samples was reacted in the presence of excess fluorine gas for 12 h in a Ni reaction vessel heated to 250°C. The SF6 generated by the reaction was first purified by removing non-condensable by-products of the reaction by cryo-separation at −120°C. A second purification was carried out by passing the SF_6_ through a GC column with ultrapure He as the carrier gas at a rate of 20 mL min^−1^. SF_6_ was isolated from residual contaminants and the carrier gas by trapping on a −192°C cold trap as the carrier gas was pumped out. The isotopic composition of the purified SF_6_ was determined on a dual inlet isotope ratio mass spectrometer in the Stable Isotope Laboratory of the Earth and Planetary Sciences Department at McGill University.

### Isotope notation

Isotopic compositions are reported using the delta notation δ^3*i*^S = (^3*i*^R_sample_/^3*i*^R_V-CDT_ − 1) · 1000; where ^3*i*^R=^3*i*^S/^32^S, *i* is 3 or 4 and V-CDT refers to the Vienna-Canon Diablo Troilite (V-CDT) international reference scale. On the V-CDT scale, the δ^34^S value of the Ag_2_S reference material, IAEA-S-1, is defined as −0.3‰ (31). The uncertainty on the measured δ^34^S values is less than ±0.2‰. Capital delta notation (Δ) is used to report deviations among the fractionation relationships of ^33^S−^32^S and ^34^S−^32^S ratios: Δ^33^S = δ^33^S − 1000 · ((1 + δ^34^S/1000)^0.515^ − 1) (32,33). We assume the Δ^33^S value of IAEA-S-1 is 0.094‰ V-CDT. The uncertainty on the measured Δ^33^S values is less than ±0.01‰. Fractionation factors (^3*i*^α) between reactant (r) and product (p) are given by ^3*i*^α = ^3*i*^R_p_/^3*i*^R_r_; and ^3*i*^R = 1 + δ^3*i*^S/1000. Herein these are expressed as isotopic enrichment factors in ^3*i*^ε notation, where ^3*i*^ε (‰) = (^3*i*^α − 1) · 1000. The fractionation factors for the heavy isotopologues are related by: ^33^λ = ln ^33^α/ ln ^34^α; where ^33^α is the fractionation factor for ^33^S−^32^S ratios and ^34^α is the fractionation factor for ^34^S−^32^S ratios.

### In situ sulfate reduction rate

Dissimilatory sulfate reduction rates were determined using radiotracer ^35^SO_4_^2-^ (34). From the Outlet and Channel-3 station, six 5 cm replicate sediment cores were taken using cut-tip PE syringes. Immediately following collection, each core was capped with a butyl stopper, wrapped in electrical tape, and 120 μl of ^35^SO_4_^2-^ (1480 kBq) was injected along the central vertical axis of each core. Syringe cores were returned to the holes from which they were derived, and each core was incubated *in situ*. Expecting lower SRRs than are typically found in coastal sediments (~20 nmol SO_4_^2−^ cm^−3^ d^−1^, where sulfate concentration is 20 μmol cm^−3^), higher amounts of tracer and longer incubations were employed (34). Microbial sulfate reduction was terminated at time points 0, 24 and 72 h from each station by expelling sediment core material into 35 mL 20% wt/wt ZnAce, precipitating H_2_^35^S as Zn^35^S. Samples were stored and transported frozen. Subsamples of the ZnAce from fixed samples were taken to determine ^35^S_sulfate_. Reduced sulfur and unreacted sulfate were separated by hot acidic distillation in the presence of 1 M Cr^2+^ (CrCl_3_(OH_2_)_6_). Radioactive sulfide was captured into ZnAce. Precipitated Zn^35^S was combined in 20 mL vials with scintillation cocktail. Radioactivity of all samples was quantified on a liquid scintillation counter. The fraction (*F*) of sulfate reduced was determined by the ratio of total reducible inorganic sulfur (TRIS) activity to total activity as *F* = A_TRIS_ / A_*SO42−*_ + A_TRIS_. Sulfate reduction rate (SRR) was calculated as *SRR* = *F* · [*SO_4_^2−^*] · 1.06 · *ϕ* / *t*, where [SO_4_^2−^] is the pore water sulfate concentration, ϕ is porosity, and t is incubation time. The factor 1.06 is the estimated isotope fractionation between ^32^S and ^35^S during bacterial sulfate reduction (34).

### DNA extraction, amplification and sequencing

DNA was extracted from 0.5 g sediments. Individually barcoded 16S rRNA gene and *dsrB* amplicons from each DNA samples were prepared using a 2-step PCR approach according to recently established workflows (25,35). Negative controls of the barcoding procedure were performed with ddH_2_O as a template and negative control amplicons were included in all further processing steps. Pooled libraries were sequenced on an Illumina MiSeq system (35). Sequenced datasets are available in the NCBI Sequence Read Archive; Project ID PRJNA512289. Further descriptions of DNA extraction, amplification, and purification are given in Supplemental Materials: Methods and Table S2.

### Bioinformatics, OTU classification and diversity calculations

16S rRNA gene and dsrB raw sequencing reads were demultiplexed and quality filtered as previously described in Herbold *et al*., 2015 and Pelikan *et al*., 2016, respectively. OTU clustering of 16S rRNA gene amplicons was performed in USEARCH (36) in a two step process with OTU clustering (-cluster_otus) at 97 %. These OTUs were classified using the Ribosomal Database Project naïve Bayesian classifier (37). OTU clustering of *dsrB* amplicons and taxonomic classification of OTUs was performed as previously described (25). OTU tables and classification tables were imported into the R software environment (R core team, 2014) and analyzed there using native functions and the sequence data processing software package phyloseq (39). First both datasets were filtered for samples with more than 100 reads. Then relative abundances were calculated and OTUs with the highest relative abundance in the negative controls were removed from the datasets. Additionally, samples with a Bray-Curtis distance of less than 0.8 to the negative control were removed from the datasets. Only samples present in 16S rRNA gene and dsrB dataset were kept for further analyses. The “clean” OTU tables were then rarefied at the smallest library size. Alpha diversity indices for species richness (Chao1) and richness and evenness (Shannon) were calculated for both 16S and *dsrB* for each sampling station (40,41). PCoA plots of environmental influences on microbial beta diversity were made from Bray Curtis dissimilarity matrices using the R community ecology package ‘vegan’ (42).

### Modeling the environmental cell-specific sulfate reduction rate-isotope fractionation relationship

We applied the most recent mechanistic model of sulfur isotopic fractionation by SRM (43) to our environmental system to evaluate whether the general parameters of the model, informed by cultured isolates, can accurately predict fractionations expressed *in situ*. The model relies on measured environmental parameters (including concentrations of sulfate and sulfide, and temperature), equilibrium and kinetic fractionation factors, and the thermodynamics that determine the reversibility of each step of the reduction pathway (43). Each of these parameters has been empirically evaluated for at least one, and often several SRM. To evaluate the potential for species-weighted sulfate reduction pathway enzyme kinetic parameters deviating from the values implemented in the model, each of these enzymatic parameters (V_max_, K_s_, K_p_), for each enzymatic step was varied over several orders of magnitude to assess the extent to which each would need to deviate from the model default in order to result in the net fractionation observed. By evaluating the modeled relationship between environmental sulfate reduction and the preserved signal of isotopic fractionation, we take a first step in exploring whether the conundrum of genomic variability resulting in fractionation variability between isolate cultures (12) is relevant to the net fractionation of sulfur by a community of organisms.

## 3. Results

### Sediment and pore water chemistry

Moving downstream from the anoxic outlet sediments, channel sediments trended towards more oxygenated, warmer, higher pH, less negative redox potential, lower concentration of dissolved sulfide and a greater concentration of dissolved organic carbon (Table S3). Sulfide values below the detection limits of our assay (16 μM), were treated as having concentration 8±8 μM.

### Sulfur isotopic composition of water and sediments

The majority of sulfur in GH4 sediments was chromium-reducible sulfur (CRS; Fig. S1). Sulfur isotope values (δ^33^S and δ^34^S) derived from mineral gypsum from Gypsum Hill, and GH4 spring water and sediments are listed in Supplemental Materials, Table S4. Soluble sulfate δ^34^S values were 17.6 ‰ for GH4 spring water, 18.6 and 19.4 ‰ for gypsum, and ranged between 16.7 and 20.2 ‰ in GH4 sediment pore water. Insoluble elemental sulfur (S^0^) was not recovered from spring water. GH4 sediment δ^34^S^0^ values ranged between -29.0 and -25.5 ‰. Acid volatile sulfides (AVS) were in insufficient concentration (< 40 μg S g wet sediment^-1^) to recover from several sediment samples (Fig. S1); δ^34^S_AVS_ values were -29.9 ‰ for GH4 spring water and ranged between -26.2 and -24.3 ‰ in GH4 sediments; CRS δ^34^S values were between -27.9 and -22.0 ‰. Δ^33^S values reflected the phase of sulfur, ranging from 0.1 to 0.3 ‰ for soluble sulfate and from 0.10 to 0.15 ‰ for elemental and reduced forms (Table S4). No increasing δ^34^S_sulfate_ trend with distance from outlet was observed (Supplemental Materials Fig. S2), suggesting that spring water sulfate isotopic depletion was inconsequential to downstream source sulfate.

### Sedimentary record of microbial sulfur isotopic fractionation

In order to constrain the fate of metabolic waste sulfide and the potential for additional isotopic fractionations associated with secondary metabolisms, we take as possible fates: 1) remaining soluble in sediment pore water, subject to export, 2) reaction with metals and precipitation as metal sulfides, 3) biological or abiotic oxidation (Supplemental Materials, Fig. S3). The flux (φ) to each pool is unknown, and so we plot outlier values, assuming φ to each pool is 1 (totality), using the measured δ^34^S values separately for each recovered pool of AVS, CRS and S^0^ per sampling station (Fig. S1). By considering each case of φ=1, the fractionation to that pool from sulfate is equal to the fractionation between source sulfate and the sulfide produced by microbial sulfate reduction. In cases where each of these pools was recoverable, this generates a triangle of boundary fractionation values resulting from microbial sulfate reduction. The range of apparent isotopic fractionations of ^34^S (^34^ε_sulfide-sulfate_) from the sediment samples were plotted against the ratio of ^34^S to ^33^S fractionation (^33^λ_sulfide-sulfate_) and found to fall within the area bounded by in vitro experimental data derived exclusively from SRM isolates (Fig. 3).

**Fig. 3.**
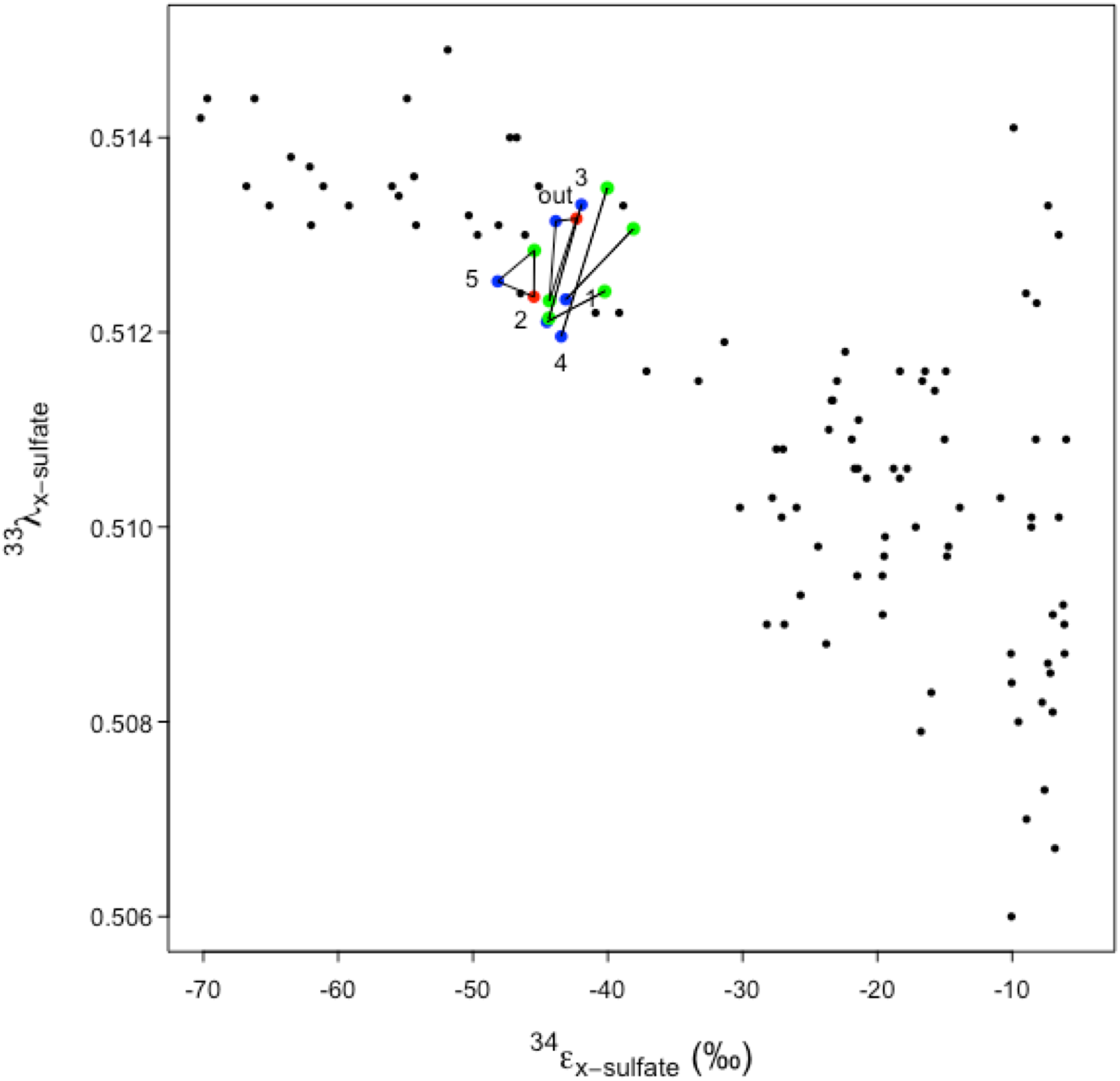
Microbial sulfate reduction boundary fractionations. Triangle apices represent ^33^λ, ^34^ε fractionation values if flux of AVS to product x is complete. x is elemental sulfur (S^0^; blue data points), chromium reducible sulfur (CRS; green data points), or hydrogen sulfide (H_2_S; red data points). Background points are assembled from literature of cultured sulfate reducers and all values are ^34^ε_sulfide-sulfate_. (Sim *et al*. 2011a, Sim *et al*. 2011b, Leavitt *et al*. 2013, Pellerin *et al*. 2015). Isotopic notation (^33^λ, ^34^ε) is described in Methods. Sampling station (out=Outlet, numbers indicate channel stations) is indicated nearest to x=S_0_ value. ^33^λ, values associated with microbial sulfur disproportionation typically exceed 0.515 (Johnston et al. 2005).

### In situ rates of sulfate reduction

*In situ* SRRs were measured from the outlet and a single channel sampling station 8 m downstream (Channel-3). Replicate one-day incubation periods yielded SRRs of 0.05 and 0.08 x 10^-9^ moles cm^-3^ d^-1^ from the outlet station and 0.24 and 0.27 x 10^-9^ moles cm^-3^ d^-1^ from the channel station. Replicate three-day incubations yielded SRRs of 0.12 and 0.21 x 10^-9^ moles cm^-3^ d^-1^ from the outlet station and 1.32 and 1.91 x 10^-9^ moles cm^-3^ d^-1^ from the channel station. Calculated fractions of reduced sulfate from replicate control outlet and channel station incubations that were fixed immediately following addition of ^35^S tracer were 1 to 2 orders of magnitude lower than fractions reduced in samples incubated for one or three days. Minimum rates of *in situ* csSRR are plotted by dividing measured bulk *in situ* sulfate reduction rate by the total number of cells in the same sample volume (see Colangelo-Lillis *et al*. 2017 for reporting of cell counts from parallel sample collection). These are plotted against the range of net preserved signals of S isotope fractionation from the Outlet and Channel-3 (Fig. 4). On the same plot are measurements from a suite of SRM grown in vitro. For the rates of reduction observed, fractionation was lower than would be predicted from SRM grown under controlled conditions.

**Fig. 4.**
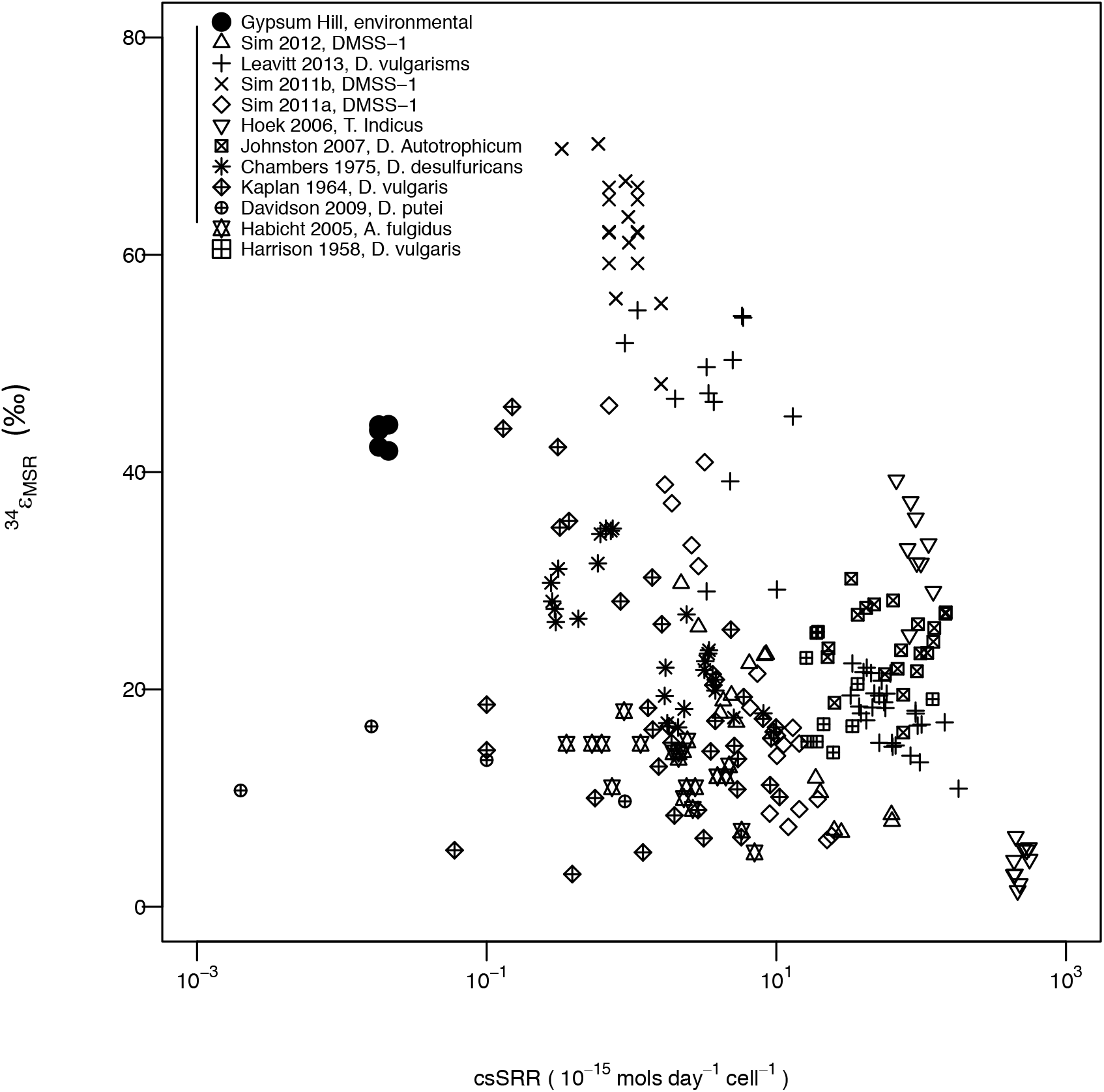
Gypsum Hill Spring sediment and cultured isolates cell specific sulfate reduction rates (csSRR) and net isotopic fractionations (^34^ε). csSRR for GH sediments are calculated from total cell counts, fractionation value ranges are from Outlet and Channel 3 stations and reflect range of possible fractionations including all potential fluxes of sulfide (filled circles). All other measurements represent sulfate fractionation associated with variable csSRR from a pure culture studies: First author, year of study and organism are indicated by marker symbol. In keeping with other literature, isotopic enrichment fractionation ^34^**ε**_sulfide-sulfate_ is presented here as ^x^**ε** = 1000*(1−^x^**α**).

### Relative abundance and diversity of 16S rRNA gene and dsrB

A total of 357,494 high quality 16S rRNA gene sequences were obtained, with an average of 44,687 sequences per sample replicate (n=8). For one replicate from each of the Outlet and Channel-4 stations too few reads were recovered or the community composition was too close to the negative PCR control indicating contamination and only a single replicate was used for analyses. A high Good’s coverage (homologous coverage) of 98-99% (Table S5) of the libraries shows that most of the gene diversity in the amplicons was represented by the recovered sequences. Overall analysis of operational taxonomic units (OTUs) at the approximate species-level (97% sequence similarity) showed that across all samples the sediments were moderately diverse (Chao1 208-549, Shannon 2.22-3.59; Table S5) and dominated by the classes *Gammaproteobacteria* (18-66% of sequences), *Deltaproteobacteria* (4-14%) and *Clostridia* (2-26%; Fig. 5A). Within the known SRM taxa, the majority of sequences belonged to families *Desulfuromonadaceae* (23-62% of *Deltaproteobacteria* sequences), *Desulfobulbaceae* (19-42%) and *Desulfobacteraceae* (3-44%; Fig. 5B). Remaining *Deltaproteobacteria* families each made up less than 2% of the class. *Desulfuromonadaceae* exhibited a marked increase in abundance with distance from outlet, while *Desulfobacteraceae* exhibited a marked decrease along the same gradient.

**Fig. 5.**
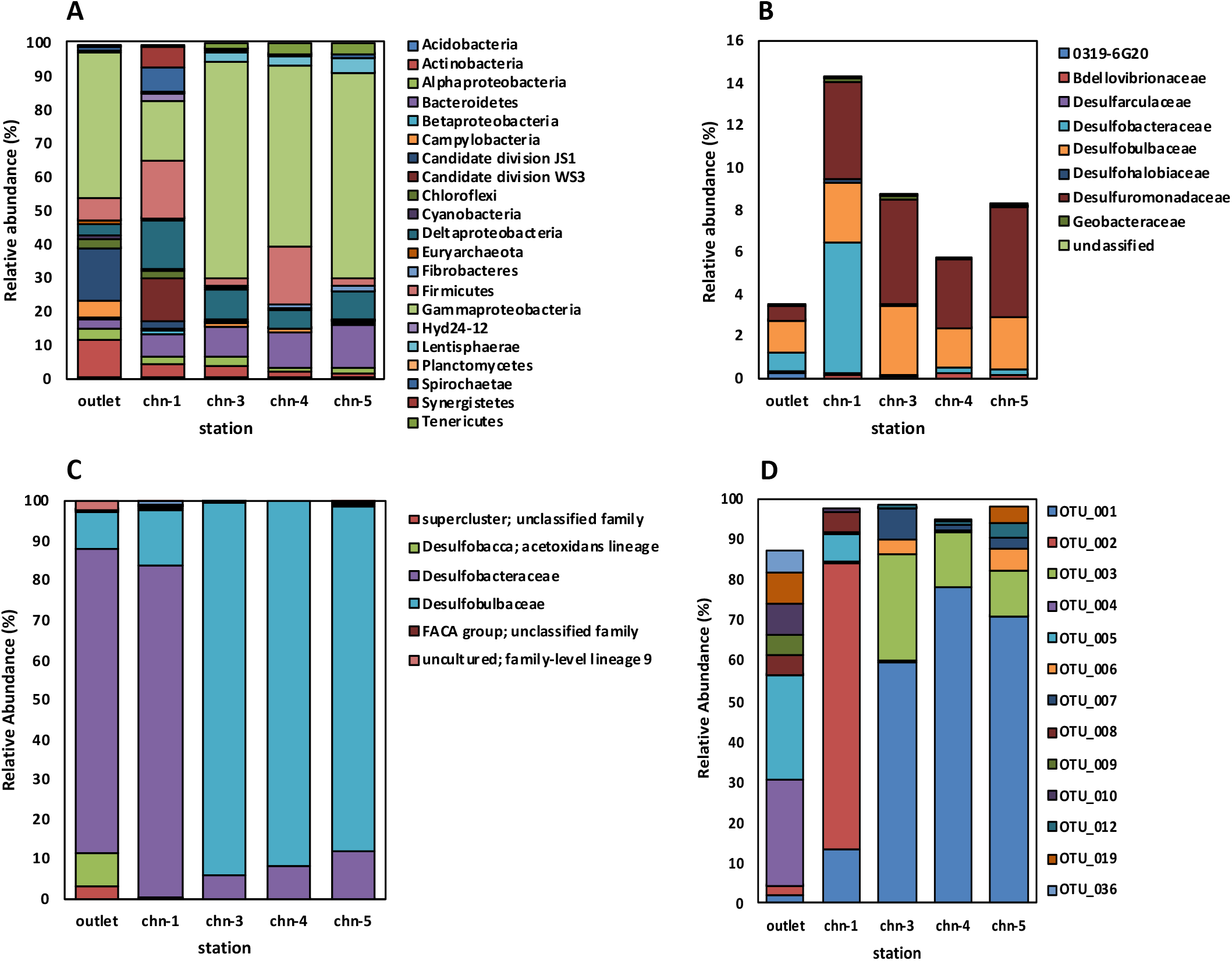
16S rRNA and *dsrB* gene identity and relative abundance. In each plot only taxa at > 1% relative abundance are shown. A) 16S rRNA; phyla B) 16S rRNA; deltaproteobacterial families. C) *dsrB*; described taxa and uncultured, family-level lineages. D) *dsrB*; species-level OTUs (taxonomy not assigned, plot only illustrates relative OTU-level diversity distributions).

A total of 140,441 high quality *dsrB* sequences were obtained, with an average of 17,555 sequences per sample replicate (n=8). Overall analysis of OTUs at 99% sequence similarity (25) showed lower diversity compared to 16S rRNA-OTUs (Chao1 15-52, Shannon 0.80-2.17; Table S5) and that *Desulfobacteraceae* (6-84%) and *Desulfobulbaceae* (9-94%; Fig. 5C) dominated. The relative abundance of these *dsrB* sequence types showed notable trends in the spring sediments; *Desulfobacteraceae* dominated the outlet and upstream channel sediments (77-84%) and *Desulfobulbaceae* dominated downstream channel sediments (87-94%). Remaining sequences were most closely related to families that composed less than 2% of all *dsrB* sequences averaged across all stations. Sequences related to the *Desulfobacca acetoxidans* lineage were the only other *dsrB* sequence type to exceed 5% at of any individual station (Outlet; 8.7%). *dsrB* OTU distributions were more even for the Outlet station compared to all Channel stations (Fig. 5D).

As a first step in evaluating the role of a SRM community on the net fractionation of sulfur isotopes in the environment, we queried a number of measured environmental factors at each sediment sampling site, for correlation with alpha diversity indices of species richness (Chao1) and both richness and evenness (Shannon) for both the 16S rRNA gene and *dsrB* (Fig. 6). Correlations were strongest between temperature and *dsrB* Chao1 diversity (R^2^=0.93, p=0.01), oxygen and *dsrB* Chao1 diversity (R^2^=0.81, p=0.04), and pH and 16S rRNA Shannon diversity (R^2^=0.88, p=0.02)(Fig. S4); all other correlation R^2^ values were less than 0.65 and p values were greater than 0.1. Similarly, the same metrics of alpha diversity were plotted with net fractionation. No significant correlations were measured (R^2^≤0.3, p≥0.3)(Table S6).

**Fig. 6.**
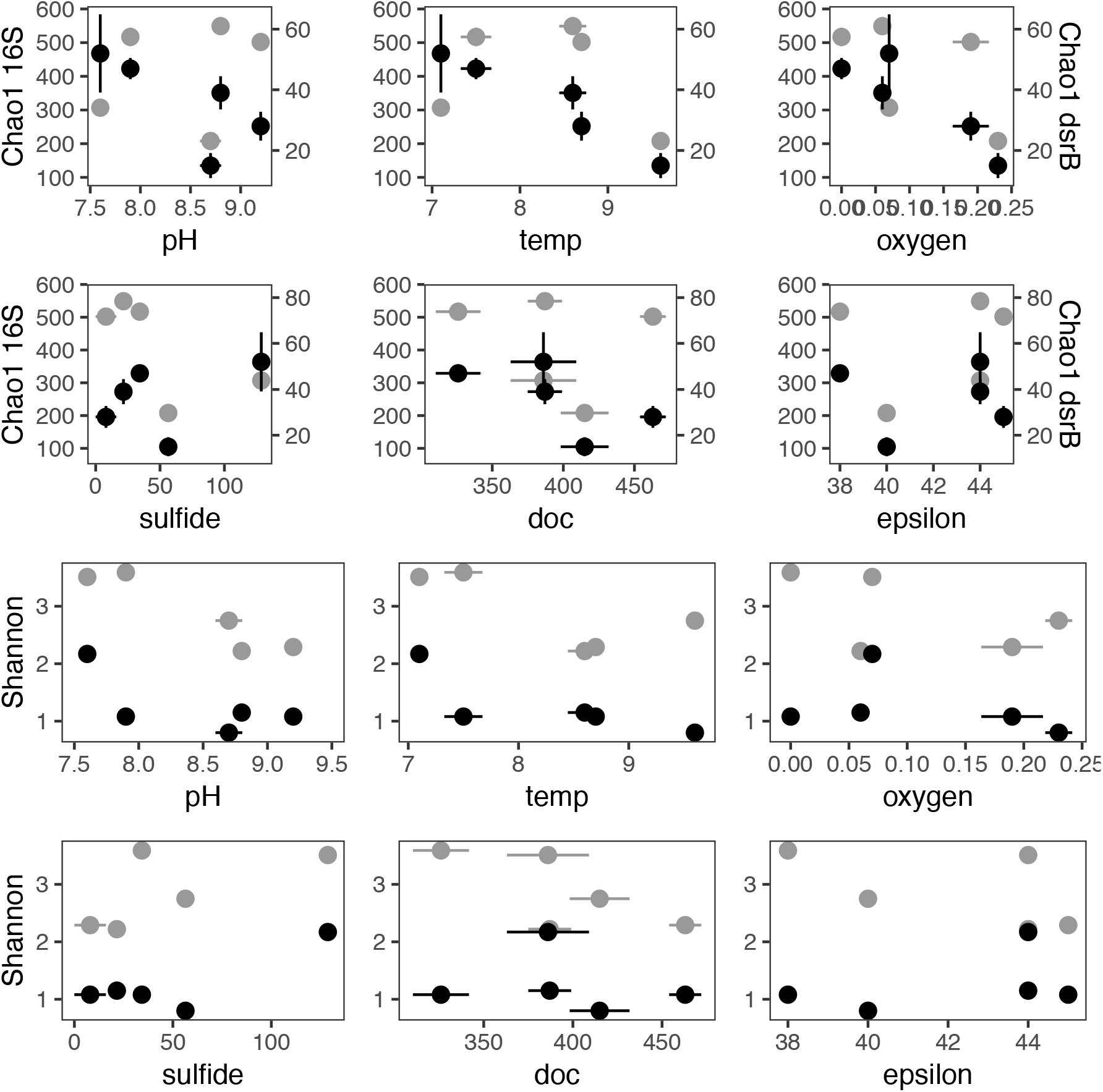
16S rRNA and *dsrB* gene diversity correlations with environmental factors and sulfur isotope fractionation. Chao1 (first and second row) and Shannon (third and forth row) diversity metrics from Outlet and each Channel station are plotted against environmental parameters: pH, temperature (°C), dissolved oxygen (**μ**M), dissolved sulfide (**μ**M) and dissolved organic carbon (doc, **μ**M); and net sulfur isotopic fractionation (epsilon, ‰). 16S rRNA Chao1 diversity (grey circles) values correspond to primary (left-hand) y-axes of upper two rows, *dsrB* Chao1 diversity (black circles) values correspond to secondary (right-hand) y-axes of upper two rows. 16S rRNA and *dsrB* Shannon diversity values are plotted on the same scale. Error bars indicate standard deviations of triplicate measurements of environmental parameters and isotope *δ*^34^S measurements, and standard errors of diversity indices calculations.

### Modeling the environmental cell-specific sulfate reduction rate-isotope fractionation relationship

Under the measured environmental parameters from GH4, net fractionation values (70 ‰) from a recent bioisotopic model of sulfate reduction (43) were much greater than observed (40-45 ‰). By adjusting the number of cells contributing to the bulk csSRR, creating a category of ‘SRM+active’, csSRRs predicted to generate the fractionations (^34^ε) measured from GH4 implied ≈50-100 times greater single cell activity than estimated from bulk cell counts. In addition, no individual enzymatic kinetic parameter could be varied in the model by less than one order of magnitude to generate the observed fractionation. Varying kinetic parameters associated with activated sulfate reduction by adenosine 5’ phosphosulfate reductase had the greatest impact on matching the cultured isolate based model towards predicting observed fractionation (Fig. 7 and Fig. S5).

**Fig. 7.**
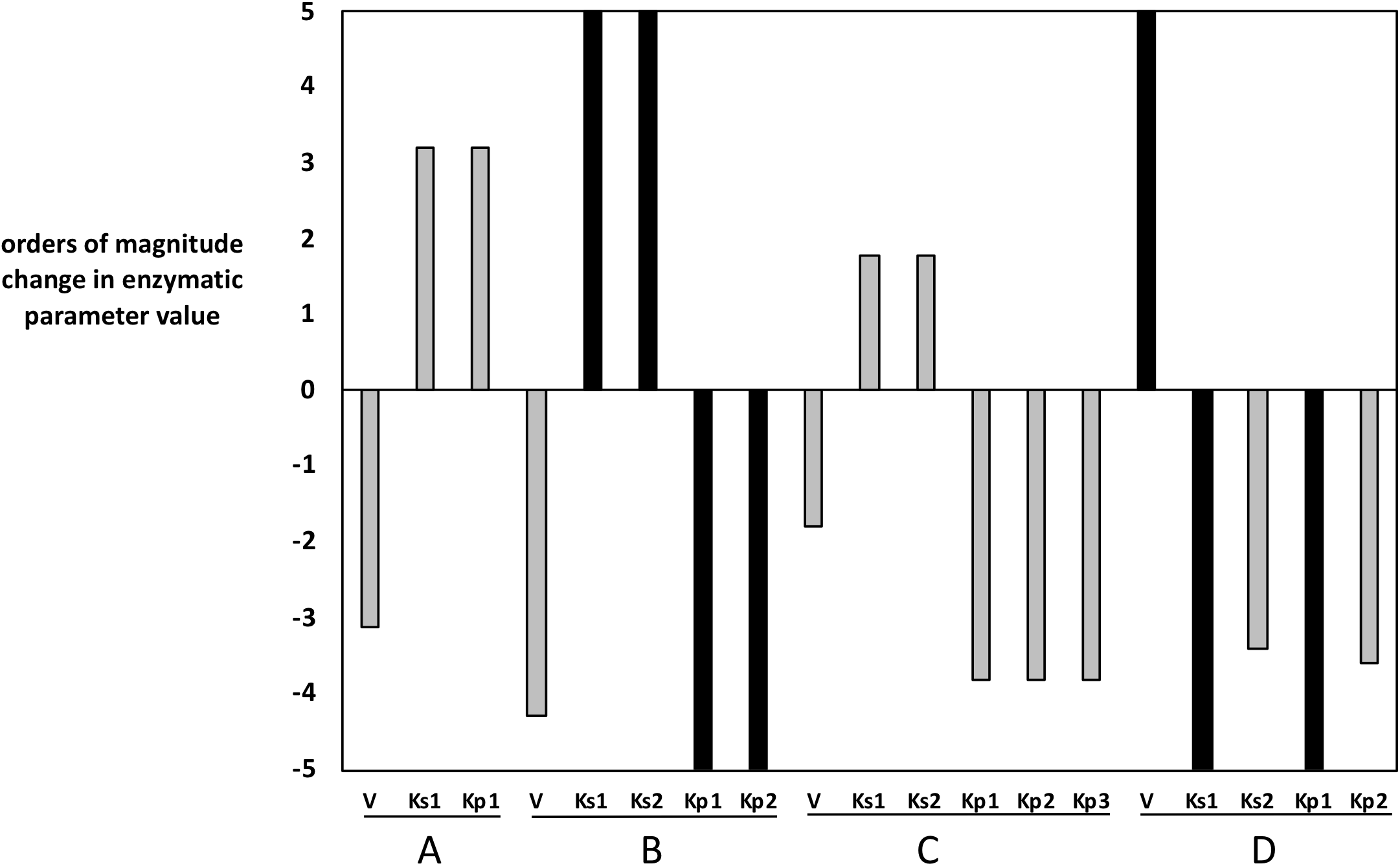
Influence of changing sulfate respiration pathway enzymatic kinetic parameters (V-max rate ; K- half saturation constant) on modeled csSRR-fractionation (^34^ε_sulfate-sulfide_) relationship. Bar plot values indicate value of order of magnitude change required of each individual enzymatic parameter required to match predicted fractionation to that observed. Values exceeding 5 are plotted at 5. Bars colored black indicate parameters for which no value will result in predicted fractionation matching measured. Uppercase letters designate steps in the sulfate reduction pathway: A- sulfate uptake, B- sulfate activation, C-sulfate reduction, D-sulfite reduction. Lowercase subscripts indicate substrate or product: A. K_s1_&K_p1_: sulfate; B. K_s1_: sulfate, K_s2_: ATP, K_p1_: APS, Kp_2_: PPi; C. K_s1_: APS, K_s2_: MK_reduced_, K_p1_: MK_oxidized_, K_p2_:sulfite, K_p3_: AMP; D. K_s1_: sulfite, K_s2_: MK_reduced_, K_p1_: sulfide, K_p2_: MK_oxidized_.

## 4. Discussion

The goals of this study were to evaluate whether a greater SRM species diversity would be reflected in the magnitude of isotopic fractionation of sulfur isotopes, and also to compare fractionation by a well characterized community *in situ*, to the fractionation observed from cultured isolates. We described the sulfate reducing microbial community and the geochemical parameters expected to influence that community, along a hypersaline Arctic spring sediment transect. We analyzed the isotopic signature of sulfur fractionation along the same transect. Finally, we compared our findings to those predicted by an isotope fractionation model informed by the geochemistry we measured. In doing so, we apply a foundation of sulfur isotope fractionation measurements and models made from cultured isolates to an environmental system.

### Fractionation signal consistent with sulfate reduction

In the case of sediments from each of the GH4 sampling stations, measured ^34^ε_sulfide-sulfate_ and ^33^λ_sulfide-sulfate_ values were consistent with those measured in cultures of microbes solely reducing sulfate and fall outside the ^33^λ range in which microbial sulfur disproportionation can be empirically evaluated to have played a role in further metabolic processing of sulfides oxidized to sulfur (44). This assessment of fractionation resulting solely from MSR allows these results to be interpreted within the framework illustrated in Fig. 1, without additionally accounting for the influence of reduced sulfide cycling by biological reoxidation. Additionally, we can evaluate our findings against those predicted by a thermodynamics-based model of sulfur isotope fractionation associated with dissimilatory sulfate reduction. Calculated fractionation values close to 40 also justify the ^35^S fractionation factor of 1.06 in calculating sulfate reduction rate (c.f. Røy *et al*., 2014), as sulfur isotope fractionation resulting from MSR is expected to be mass dependent.

### Alpha diversity metrics correlate with environmental parameters but not isotopic fractionation

Each of the environmental factors queried for correlation with gene alpha diversity metrics has been demonstrated to play a role in shaping microbial communities in other environments; e.g. temperature: (45), pH: (46), oxygen: (47), sulfide: (48), and organic carbon: (49). Within the set of environmental parameters measured in this study, alpha diversity (Shannon) of species-level 16S rRNA gene-OTUs was best correlated with pH (Fig. 6 and Fig. S4), while species-level *dsrB* OTU richness (Chao1) was best correlated with the small differences in oxygen concentration and temperature between stations. While causality is not evaluated by this study, the relationship between 16S rRNA gene diversity with pH is consistent with numerous other studies that have explored this relationship (50,51), though notably the directionality of the relationship appears related to the absolute pH, with greatest diversity found at pH 7 and lower diversity at both higher and lower pH. The influence exerted by pH may be tied to its control on nutrient availability. That *dsrB* diversity is associated with a different set of environmental variables is not a novel finding. A similar study of 16S rRNA and *dsr* gene biogeography, across a much larger physical scale of similar environments (52) found that while 16S rRNA gene diversity correlated to environmental parameters, *dsrB* diversity did not. *dsrB* amplicon diversity responding differentially here, indicates unique selection on the subset of the microbial community that carries this functional gene. A decreasing abundance of unique *dsrB* OTUs with increasing oxygen is consistent with the anaerobic nature of most SRM and that oxygen tolerance by organisms with this metabolism differs (53,54), such that at lower oxygen concentration there may be a greater range of diversity in the genes of this pathway by nature of persistence of SRM with lower oxygen tolerance. The inverse relationship between unique *dsrB*-OTU richness and the small temperature variations in the spring is less clear. Notably there is a trend towards decreasing evenness (as measured with Shannon diversity), and it may be that small temperature increases allow for one SRM taxon to competitively edge out a number of others, decreasing dsrB-OTU richness. Given the differences in gene identity between sites, both genes appear to be dispersal limited over the small distance traversed by GH4, possibly due to constant unidirectional flow (Fig. S4).

A driving question of this work was whether changes in SRM community richness and evenness would correlate with net fractionation of sulfur isotopes. No correlation between any alpha diversity metric and net fractionation was observed (Fig. 6). Besides known caveats in using gene amplicon-based data for quantifying diversity (25,55–57), there are numerous biological explanations for this disconnect. One possible explanation is that increasing diversity of an environment favors organisms with greater csSRR (c.f. Eqn.1.). While this might be expected during ecological succession in an environment following a disturbance, it is not likely to be an ongoing process and so is unlikely to be important in this long-term stable environment. Considering the second premise, it is feasible that increasing diversity encompasses metabolically redundant individuals with both greater and lower csSRR than a less diverse environment, and that the absolute number of sulfur molecules metabolized multiplied by the fractionation associated with each organism results in net contributions to fractionation that are similar enough so as to be indistinguishable from the signal produced by the less diverse community (c.f. Eqn.1.). That is, the value of Eqn. 1 is very near to 0. In a similar way, despite differences in community composition and measured diversity indices (Figs. 5 and 6), an individual taxon may be dominating both sulfate reduction and the net fractionation signal observed at each sampling site. Without an assessment of the relative activities of the different SRM in each community, our data set does not allow us to evaluate this explanation, but points towards logical extensions of this work. The nature of addressing this hypothesis in an environmental setting may provide additional explanations to consider. There may exist a threshold of alpha diversity, beyond which further increases in diversity will no longer yield changes in fractionation. Above this threshold, differences in species-specific contributions to fractionation will blend to the point that individual contributions are overshadowed by environmental constraints on metabolite availability, in similar fashion to functional stability exhibited in environments with substantial taxonomic variability (58,59). If this is indeed the case, we can place lower limits on this threshold of *dsrB* Shannon diversity at a value of 0.8 and *dsrB* Chao1 species richness at a value of 15. If most natural environments fall into this category, with diversity above these thresholds (e.g. Jochum *et al*., 2017), understanding the specific mechanisms of fractionation within taxa is less important to determining net fractionation and interpreting the sulfur isotope rock record than are environmental conditions. This further justifies examining the relationship between diversity and net fractionation *in vitro*, on artificial communities of small sizes, that may be representative of the functional diversity present when sulfate reduction appeared in geological time. While future studies will benefit from a larger range of samples with varying alpha diversities, our data still gives a first indication that net sulfur isotope fractionation is not affected by microbial composition above a certain diversity threshold.

### GH4 sulfur fractionation values were lower than model predictions

The inverse relationship between csSRR and observed fractionation of sulfur isotopes by SRM is among the most enduring of explanations for variation in fractionation signals (1,4–6,10,61–64). It has been proposed that environmental factors influence fractionation only to the extent that they influence csSRR, and this relationship underlies the interpretation of paleoenvironments from sulfur isotopes in the rock record, based on manipulated cultures of SRM. For the minimum csSRRs observed, fractionation was lower than would be predicted from SRM grown under controlled conditions (Fig. 4). Actual csSRR of the active cell population is expected to be significantly higher, as only a fraction of the microbial community is capable of sulfate reduction, and of those, activity is likely heterogeneous. Finding that per cell activity would need to be much greater than calculated from bulk cell counts in order to match the observed fractionation with those seen in culture is consistent with the finding that only 3.5% and 8.7% of the microbes from each station are *Deltaproteobacteria*, a first order approximation for the abundance of SRM, and validated by the taxonomic assignment of sequenced *dsrB* genes largely to deltaproteobacterial families with bona fide SRM. However even by assigning the bulk sulfate reduction to these putative fractions of SRM from each population, measured csSRR was still lower than expected to generate the observed signal. Comparisons with the isotopic fractionation model based on cultured SRB (Wing and Halevy, 2014) indicated measured csSRR was still ≈2 to 10 times lower than expected to generate the observed signal. The simplest explanation to resolve this discrepancy is that only ≈60% and 10% of the *Deltaproteobacteria* from Outlet and Channel-3 sediments, respectively, are actively reducing sulfate *in situ*. Remaining *Deltaproteobacteria* may be inactive, or may metabolize different substrates (i.e. fermenting organic acids, or reducing iron or elemental sulfur, as *Desulfuromonadaceae* are known to). Recalculating the csSRR using these revised numbers of actively respiring cells, the observed fractionation matches the predicted fractionation. Heterogeneous activity within isolate cultures is well documented (65,66). Though a distribution of activity within functional groups is expected *in situ*, the methods to evaluate this activity on environmental *in situ* samples are only starting to be developed (e.g. Hatzenpichler *et al*., 2016; Berry *et al*., 2015). The influence of heterogeneous activity on apparent csSRR has substantial implications for both interpretation of calculations of per cell reduction rates and fractionation signals preserved in the rock record. Preserved fractionation signals are consistently interpreted as being generated by a population of SRB reducing sulfate at a single rate and do not consider the possibility that specific taxa may responsible for a disproportionate amount of both activity and fractionation.

### Enzymatic kinetic explanations for discrepancy between in vitro and in situ csSRR-fractionation

An alternative explanation for the deviation from predicted csSRR, is deviations in the physiological parameters influencing fractionation between those of model organisms and environmental populations. Notably populations of enzymes adapted to cold temperature environments exhibit higher specific activities than would be expected, to maintain catalytic rates comparable to orthologous genes in warmer temperature regimes (69). To assess this discrepancy as a possible cause for the differences between measured and modeled fractionation values, we manipulated in silico the enzymatic kinetic parameters V_max_ (the enzyme’s maximum rate) and K_m_ (the enzyme’s half saturation constant) associated with each enzyme, its substrates and products, in the sulfate reduction pathway (Fig. 7 and Fig. S5). These manipulations treat the microbial community diversity of each enzyme type as a single enzyme, but give an idea of the deviation for each parameter from those utilized in the model (and based on empirical values) that would be required to yield the fractionation values measured in GH4. Broadly to find better agreement between the csSRR-fractionation relationship observed in GH4 and that observed in culture, the maximum velocity of each reaction would need to be smaller, the saturation concentration of each enzyme’s substrate would need to be greater, and the saturation concentration of each enzyme’s product would need to be smaller. Of 19 enzymatic parameters (maximum rate V and half saturation constants K_m_ associated with each substrate), 16 required a manipulation of four orders of magnitude or greater from the empirical values employed in the model (see Wing and Halevy, 2014) in order to generate the measured fractionation values (Fig. 7 and Fig. S5).

Notably all three enzymatic parameters requiring a change of less than three orders of magnitude from the empirical value were associated with the Apr enzyme (i.e apr V_max_, K_M_APS_ and K_M_ATP_), which is responsible for the reduction of activated sulfate to sulfite. Two conclusions can be drawn from this exercise. First given the known diversity of activity within cultured isolates, and the expectation of similar and greater diversity of activity within a functional class of enzyme, we favor heterogeneous activity accounting for the difference between measured and modeled fractionations rather than enzymatic kinetic parameters at three or four orders of the magnitude of measured values. For example, the range of Apr [EC 1.8.99.2] K_M_ values for substrates adenylyl sulfate (forward reaction) and sulfite (reverse reaction) reported on the online BRENDA database span less than two orders of magnitude (Placzek *et al*., 2017; www.brenda-enzymes.org). In contrast, the range of Sat [EC 2.7.7.4] K_M_ values for substrates sulfate (forward reaction) and adenylyl sulfate (reverse reaction) span an impressive five orders of magnitude. We note that the potential for this discrepancy does exist, and might reflect the nature of cultured organisms that grow quickly in lab environments and may possess enzymes with substantially different kinetic potentials than the organisms that make up the bulk of environmental samples. Further, work comparing enzyme kinetics measured *in vitro* and *in vivo* have found differences approaching three orders of magnitude (71). However, while the heterogenous activity of cells in the environment is well established, the range and weighting of enzymatic kinetic parameters in situ are far less well explored. Second, variation in kinetic parameters associated with the Dsr enzyme complex are more resilient to influencing net fractionation through the sulfate reduction pathway than are comparable variations associated with the Apr enzyme (Fig. 7.). Drawing a parallel between sequence diversity and enzymatic kinetic diversity, we expect that only very large changes in kinetic diversity of *dsr* would impart observable changes in fractionation. If changes in *apr* kinetics have greater influence on fractionation, another iteration of testing the sulfate reduction pathway diversity-fractionation hypothesis might better interrogate *apr* rather than *dsrB*. This approach would inform whether there were differences in selection pressure between apr and dsr but would also require environmental characterization of *apr* akin to what has been performed for *dsrB* (25) in order to accurately capture its environmental diversity. Contrary to recent suggestions of a key role for Dsr in controlling isotopic fractionation, these results indicate it is a departure from reversibility of the Apr-catalyzed reaction that controls the fractionation upon departure from equilibrium due to increasing csSRR. The possibility of Apr acting as a bottleneck resulting in isotopic fractionation at low respiration rates was described by Harrison and Thode six decades ago (1) and merits *in vitro* experimental reconsideration in light of this and previous *in silico* analyses (Wing and Halevy, 2014).

### Interpreting geologic isotope signals independently of SRM diversity

Our findings support the continued interpretation of biologically sourced isotopic fractionation of sulfur independently of the diversity of the SRM community responsible for imparting that fractionation. Importantly, this relaxes a potentially confounding layer of complexity to such interpretation as no geological means to ascertain the composition of such communities exist, and molecular techniques to constrain the temporal diversification of functional groups are not yet capable of informing spatial ecology questions. While the GH4 spring SRB communities do not demonstrate a gene diversity-isotope fractionation relationship, it is still possible that such a correlation would be evident if you only look at the physiologically active sulfur-metabolizing microorganisms in the system. Additionally, this work informs what additional environments may be well suited to address the question. An ideal set of environmental sites would exhibit comparable geochemistry, very similar net rates of sulfate reduction, equal proportions of active members of the functional group, and a wide disparity in alpha diversity metrics. Such an environment may be characterized with substantial effort, but a more promising alternative is to create these conditions from artificial, controlled populations in a laboratory environment.

## Conclusion

This work allowed us to both examine the relationship between a fundamental ecological parameter- alpha diversity- and a geologically preserved signal, and also to apply a recent thermodynamics-based model of biological sulfur fractionation to a natural environment known to contain an active microbial sulfur metabolizing community. The species-specific nature of the cell-specific sulfate reduction rate-isotope fractionation relationship suggested that communities of small but varied composition would reflect that composition in the net fractionation of sulfur isotopes. Our findings did not indicate that increased diversity, either in deltaproteobacterial 16S rRNA gene, or sulfate reduction pathway gene *dsrB* correlated with greater measured fractionation. A significant finding of this work is the difficulty in making predictions regarding environmental communities, from the careful observation and manipulation of members of those communities in the laboratory. This study indicates the complexity of measuring *in situ* functionally redundant metabolisms and makes a first step towards teasing apart the relative contributions of specific taxa to the net products of that metabolism.

Testable predictions resulting from by this study include: 1) quantifying the heterogenous cell specific activity among members of functionally redundant taxa will yield greater congruence between *in vitro* and *in situ* values of biological sulfur fractionation; 2) enzymatic kinetic parameters associated with the majority of environmentally relevant sulfate reducing microbes are substantially different than those associated with cultured sulfate reducing microbes; 3) *apr* gene diversity better correlates with isotope fractionation than *dsr* gene diversity. Future work might take advantage of the many environmental studies that have already characterized metagenomic or proteomic diversity and couple those datasets to newly generated multiple sulfur isotope datasets from the same environments. A metagenomics (e.g. Brown *et al*., 2016) and/or metatranscriptomics or a stable isotope probing approach (e.g. ref. 73) could help to identify active microbes and quantify their activity.

## Supporting information

Supplemental Materials Table of Contents

## Acknowledgements

This research was supported by the Canadian Astrobiology Training Program (NSERC CREATE *371308-09*; BAW, LGW) through a PhD fellowship to JC, the Polar and Continental Shelf Program (PCSP; LGW) through logistical support in the field, NSERC Discovery grants to BAW (RGPIN-2014-06626) and LGW (RGPNS 305490-2012), the NSF Science and Technology Center for Dark Energy Biosphere Investigations through a postdoctoral fellowship to JC, the Austrian Science Fund (P25111-B22 to AL) and by the University of Colorado-Boulder (JC, BW). Chemical analyses were performed with the assistance of the lab of Yves Gelinas (GEOTOP, Concordia, Montreal). Kenneth Wasmund assisted with sediment extraction techniques. We gratefully acknowledge advising by Hans Røy regarding ^35^S radiotracer experimental set up and sulfate extraction and critical review of an earlier draft of the manuscript by Itay Halevy.

## Supplemental Information

Supplementary information accompanies this manuscript.

Author contributions
JC and BAW designed the study; JC, CP, KW, and IA made measurements and conducted the experiments; JC, CP, CH, and BAW analyzed the data; JC, CP, AL, LGW, and BAW interpreted the data and wrote the manuscript.

