## Supplemental Materials Table of Contents for "Diversity decoupled from sulfur isotope fractionation in a sulfate reducing microbial community"

***Accompanying***

Fig. S1. Mineralized sulfur fractions from Gypsum Hill Spring 4 sediments.

Fig. S2. Gypsum Hill Spring 4 sediments Δ^33^S-δ^34^S.

Fig. S3. Potential sulfide fates in Gypsum Hill Spring 4 sediments.

Fig. S4. Environmental influences on Gypsum Hill Spring 4 sediments microbial beta diversity.

Fig. S5. Influence of changing sulfate respiration pathway enzymatic kinetic parameters (V- max rate; K- half saturation constant) on modeled csSRR-fractionation (^34^ε_sulfate-sulfide_) relationship.

Table S1. Instruments and kits utilized in this study.

Table S2. 16S rRNA and *dsrB* primer sequences.

Table S3. Gypsum Hill Spring 4 physical and chemical measurements made from each sampling station.

Table S4. Gypsum Hill Spring 4 sediment δ^3x^S and Δ^33^S values.

Table S5. Gypsum Hill Spring 4 sediment 16S *rRNA* and *dsrB* gene amplicon sequencing coverage and diversity metrics.
