## Supplemental Figure 1 for "Diversity decoupled from sulfur isotope fractionation in a sulfate reducing microbial community"

**Fig. S1. Mineralized sulfur fractions from Gypsum Hill Spring sediments.** *y*-axis extraction fractions include Milli-Q water extraction of aqueous sulfates ($SO_{4,\ sol}$), methanol extraction of elemental sulfur ($S^0$), hydrochloric acid extraction of acid volatile sulfides (AVS), acidic chromium extraction of chromium reducible sulfur (CRS), Thode solution extraction of CRS-soluble sulfur ($CRS_{sn}$), and Thode solution extraction of CRS-insoluble sulfates ($SO_{4,\ insol}$).

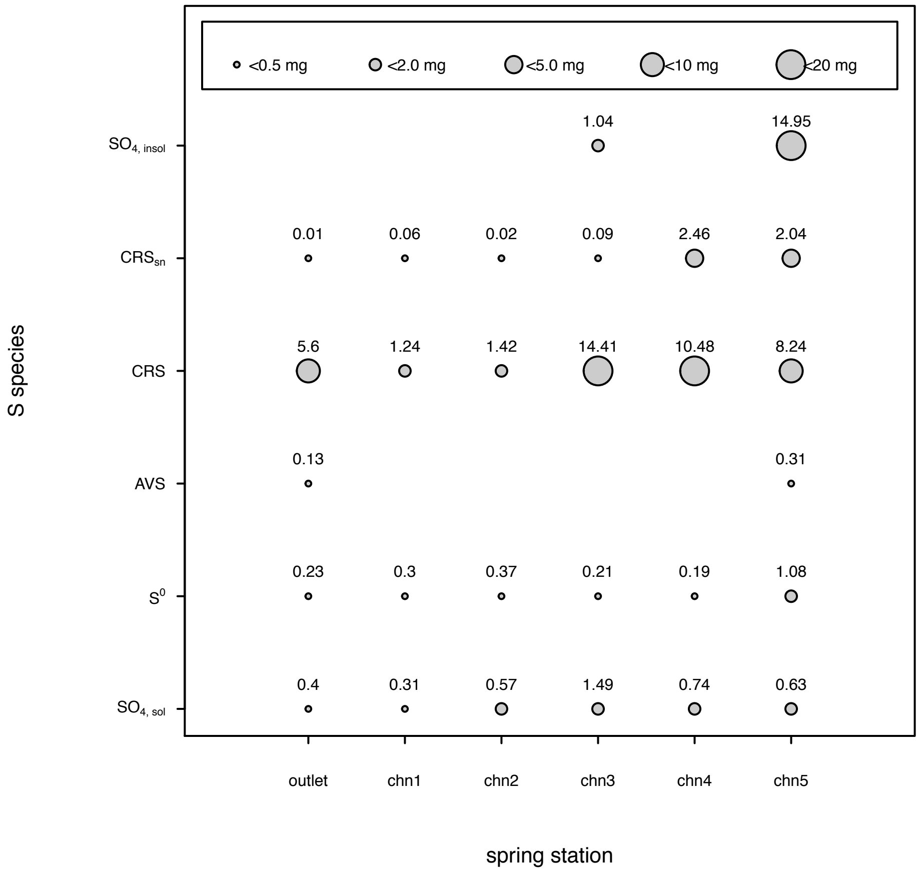
