## Supplemental Figure 2 for "Diversity decoupled from sulfur isotope fractionation in a sulfate reducing microbial community"

**Fig. S2.** Gypsum Hill Spring 4 sediments **$\Delta^{33}S$ -$\delta^{34}S$.** Symbols labeled *gyp* are from gypsum rock recovered from the spring's bank; $W_s$ are spring water samples, 'o' are Outlet samples and numbers 1 through 5 designate Channel stations 1-5.

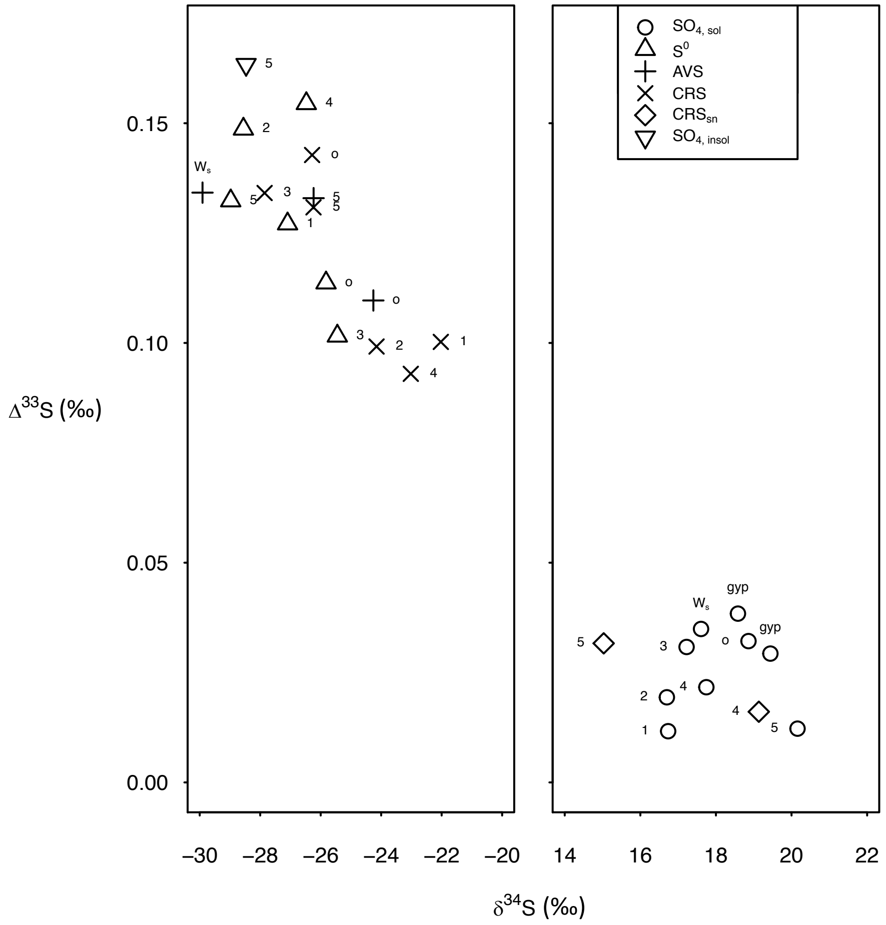
