## Supplemental Figure 3 for "Diversity decoupled from sulfur isotope fractionation in a sulfate reducing microbial community"

**Fig. S3. Potential sulfide fates in Gypsum Hill Spring 4 sediments.** $\varphi$ indicate fluxes between various S pools. AVS: acid volatile sulfur, CRS: chromium reducible sulfur.

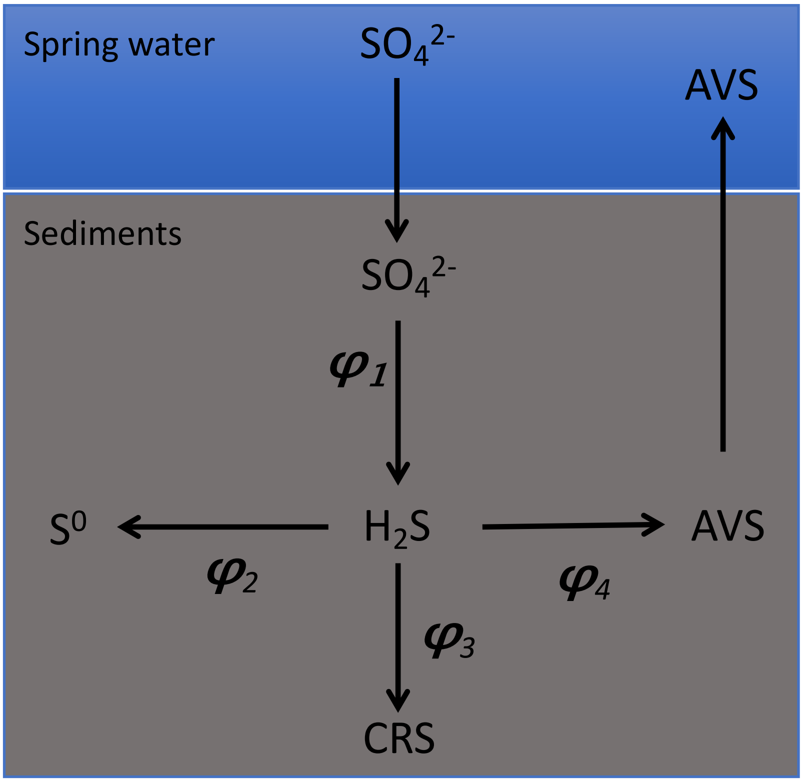


$$\varphi_1 = \varphi_{SO_4^{2-}-H_2S}$$
$$\varphi_2 = \varphi_{H_2S-S^0}$$
$$\varphi_3 = \varphi_{H_2S-CRS}$$
$$\varphi_4 = \varphi_{H_2S-AVS}$$
