## Supplemental Figure 4 for "Diversity decoupled from sulfur isotope fractionation in a sulfate reducing microbial community"

**Fig. S4. Environmental influences on Gypsum Hill Spring 4 sediments microbial beta diversity.** Principal coordinates from a principal coordinates analysis (PCoA) using Bray-Curtis dissimilarity index on equally sized re-sampled libraries are plotted against each other to summarize the microbial community compositional differences between samples. Each point represents a single sample, and the distance between points represents how compositionally different the samples are from one another. The constrained PCoA for both 16S rRNA and *dsrB* genes show the downstream channel stations are more similar to one another than to the Outlet or Chn-1 station. Oxygen, temperature, distance and pH eigenvalues are of comparable magnitude. Oxygen contributes to the variability on PC1 whereas the other environmental parameters influence both variability axes roughly equally.

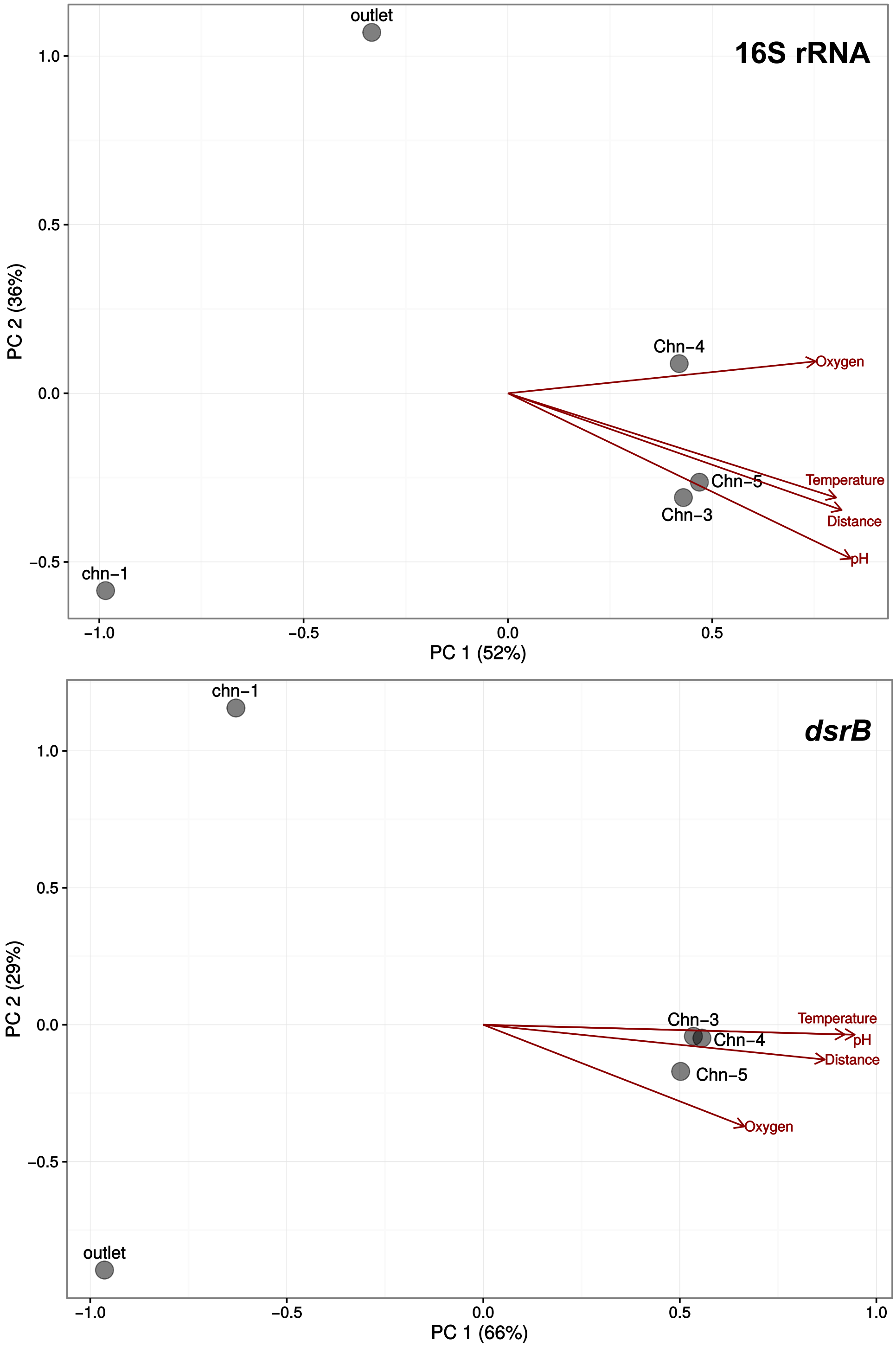
