## Supplemental Figure 5 for "Diversity decoupled from sulfur isotope fractionation in a sulfate reducing microbial community"

**Fig. S5. Influence of changing sulfate respiration pathway enzymatic kinetic parameters (V- max rate; K- half saturation constant) on modeled csSRR-fractionation ($^{34}\varepsilon_{sulfate\text{-}sulfide}$) relationship.** Average of values measured from Gypsum Hill Spring 4 sediments indicated by circle. E-value labels indicate change in relationship by changing enzyme parameter value by the indicated order of magnitude. Uppercase column letters designate steps in the sulfate reduction pathway (A- sulfate uptake, B- sulfate activation, C-sulfate reduction, D-sulfite reduction), lowercase subscripts indicate substrate or product; these are given in each plot.

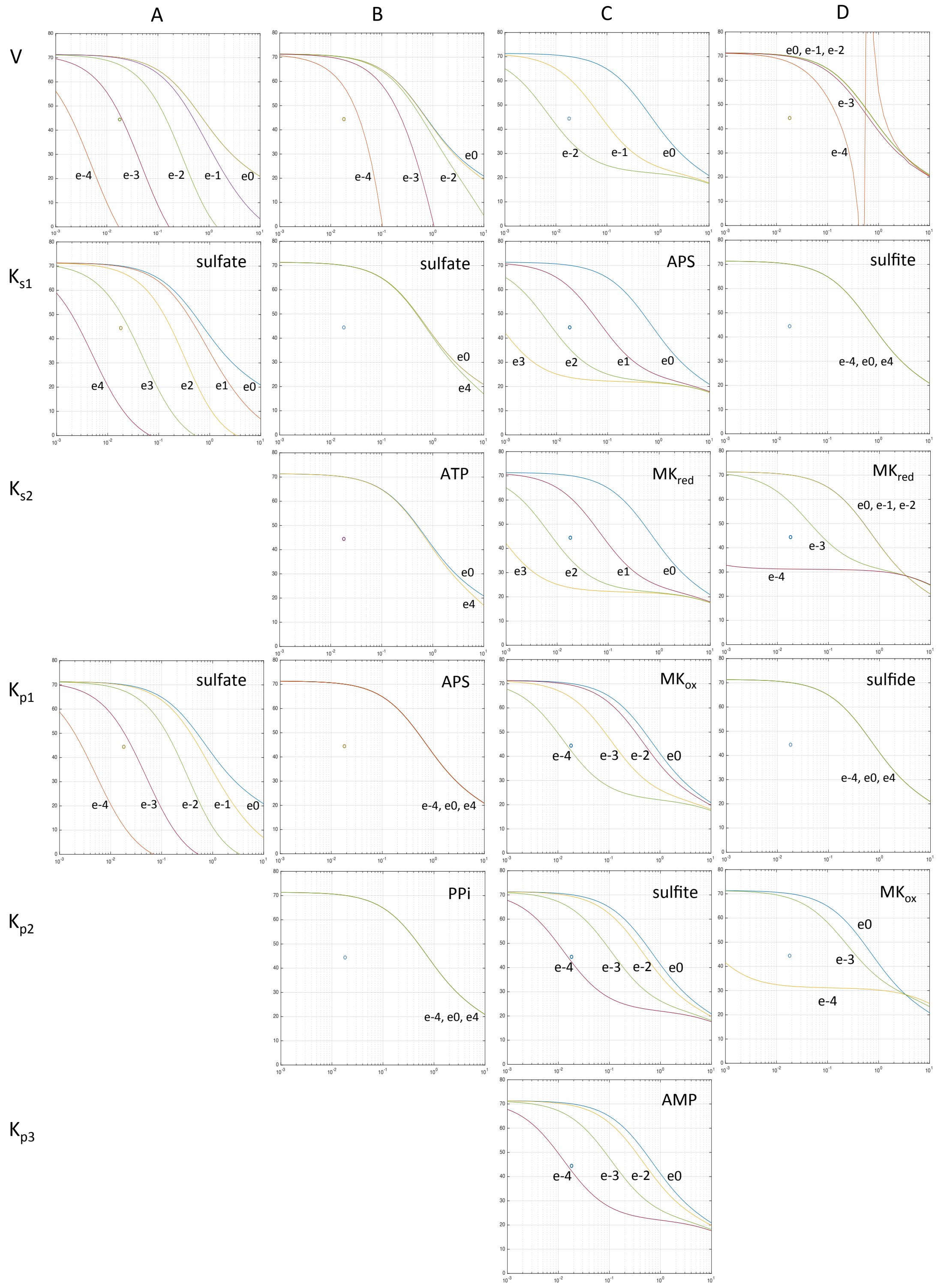
